# Assessing the impact of Brd2 depletion on chromatin compartmentalization

**DOI:** 10.1101/2024.03.02.583085

**Authors:** Advait Athreya, Liangqi Xie, Robert Tjian, Bin Zhang, Zhe J. Liu

**Affiliations:** Department of Chemistry, Massachusetts Institute of Technology, Cambridge, MA, USA; Cancer Biology and Infection Biology, Lerner Research Institute, Cleveland Clinic, Cleveland, USA; Department of Molecular and Cell Biology, Li Ka Shing Center for Biomedical and Health Sciences, CIRM Center of Excellence, University of California, Berkeley, CA, USA; Howard Hughes Medical Institute, Berkeley, CA, USA; Janelia Research Campus, Howard Hughes Medical Institute, Ashburn, VA, USA

**Author notes:** corresponding authors: Robert Tjian, Bin Zhang, Zhe J. Liu. equal contribution (Advait Athreya, Liangqi Xie).

## Abstract

Recent insights into genome organization have emphasized the importance of A/B chromatin compartments. While our previous research showed that Brd2 depletion weakens compartment boundaries and promotes A/B mixing ^1^, Hinojosa-Gonzalez et al.^2^ were unable to replicate the findings. In response, we revisited our Micro-C data and successfully replicated the original results using the default parameters in the cooltools software package. We show that, after correcting inconsistencies with the selection and phasing of the compartment profiles, the decrease in B compartment strength persists but the change in compartment identity is to a much lesser extent than originally reported. To further assess the regulatory role of Brd2, we used saddle plots to determine the strength of compartmentalization and observed a consistent decrease of compartment strength especially at B compartments upon Brd2 depletion. This study highlights the importance of selecting appropriate parameters and analytical tools for compartment analysis and carefully interpreting the results.

## Introduction

Over the past decade, advances in chromosome conformation capture technologies have revolutionized our understanding of chromatin organization, revealing the intricate three-dimensional architecture of genomes^3,4^. These technologies have uncovered a complex hierarchical structure within mammalian genomes, characterized by compartments, topologically associating domains (TADs), and loop domains, with Hi-C maps playing a crucial role in visualizing the spatial segregation of active euchromatin (A compartments) from inactive heterochromatin (B compartments). However, the molecular mechanisms driving compartmentalization remain poorly understood.

Our recent findings underscore the significant role of the bromodomain and extraterminal domain family protein Brd2 in mediating enhanced compartmentalization and the spatial mixing of accessible chromatin following cohesin depletion^1^. High-resolution Micro-C^5,6^ experiments further support that Brd2 depletion induces noticeable weakening of B compartment and A/B mixing in the presence of cohesin, aligning with its known selective association with CTCF to maintain architectural boundaries^7^. Consistent with previous reports in a different cell model^8^, we discovered that endogenous Brd2 forms complexes with CTCF independent of nucleic acids, with a pronounced enrichment at CTCF binding sites among BET family proteins in mouse embryonic stem cells^1^. These insights were garnered using established Hi-C and Micro-C analysis pipelines, specifically utilizing cooltools^9^ for A/B compartment score assessment.

Contrastingly, a recent study employing the dcHiC computational framework^10^ has reported findings that diverge from ours^2^. To address this discrepancy, we undertook a comprehensive reassessment of our data and analysis. This reevaluation demonstrated that we could reproduce our original results performed by cooltools using the default parameters. Nevertheless, a detailed examination of per-chromosome profiles uncovered discrepancies in the compartment profile selection and potential variations in statistical calculations between the two pipelines. Saddle plots and saddle strength analyses suggest that Brd2 depletion weakens compartmentalization particularly at B compartments.

## Results

### Published results from Xie et al. 2022 can be replicated using default settings in cooltools at 1Mb resolution

We processed the submitted Micro-C datasets using standard Hi-C pipelines^11,12^, and computed compartment scores for contact maps at 1Mb resolution using the *eigs_cis* function provided in the cooltools software package for Hi-C analysis^9^. We used a standard GC content track for orienting the eigenvectors and the program defaults to choose the first eigenvector without employing any selection methods. With this approach, we were able to nearly recreate the original published histogram and compartment profiles for chromosomes 11 and 17 (Fig. 3(a), Fig. 3(b), Extended Data Fig. 8(b) from Xie et al.^1^) (Figure 1a-b).

**Figure 1.**
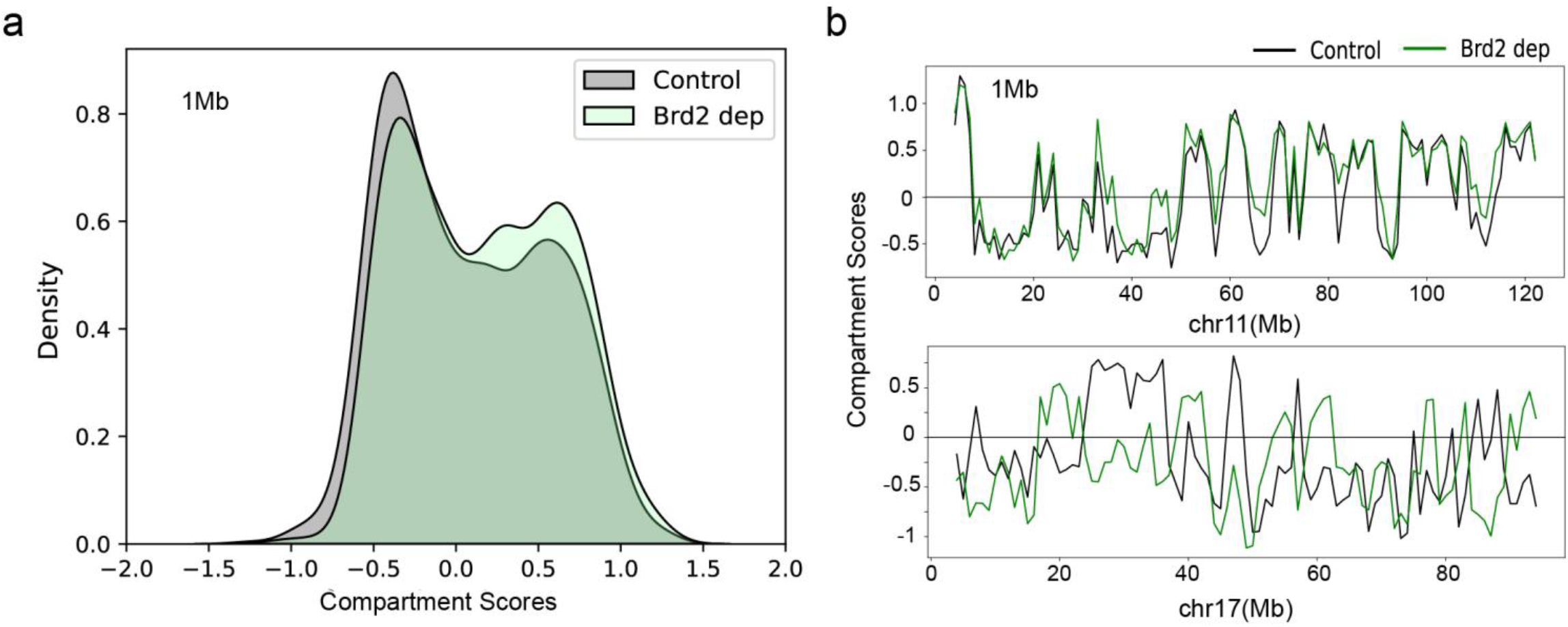
Compartment analysis performed with cooltools at 1Mb resolution yields compartment score distribution (Panel a) and chromosome-wide profiles (Panel b) closely mirrors Fig. 3(a), Fig. 3(b), Extended Data Fig. 8(b) reported by Xie et al.^1^ Only the first eigenvector was computed, and the orientation was chosen to correlate with GC content. It is worth noting that, only the reversed orientation of the compartment profile for chr17 matches the original panel, likely due to the use of a non-standard active histone mark phasing track in the original publication. The profile for chr17 appears to have poor concordance with the control in either orientation, however. Brd2 depletion profiles are only shown for replicate 2.

### The trend persists after correcting the eigenvector selection and orientation inconsistencies

As the original results appear to have had some inconsistent or inaccurate eigenvector selection and orientation, we updated the analysis by manually selecting and orienting eigenvectors using the correlation with both GC content and gene density. This approach is used in the software package dcHiC^10^ for selecting compartment profiles and is the tool employed by Hinojosa-Gonzalez et al. 2023 in their replicability study^2^. We also include both replicates for which data is available in the supplementary files. With this correction, we see that the observed trend is still present, although it is stronger in replicate 2 than in replicate 1 (Figure 2a). In addition, we observe concordant compartment score shifts from B to A at similar locations along individual chromosomes across both replicates (Figure 2b). However, the extent of changes in compartment identity, after correcting the eigenvector selection, is significantly less pronounced than initially reported.

**Figure 2.**
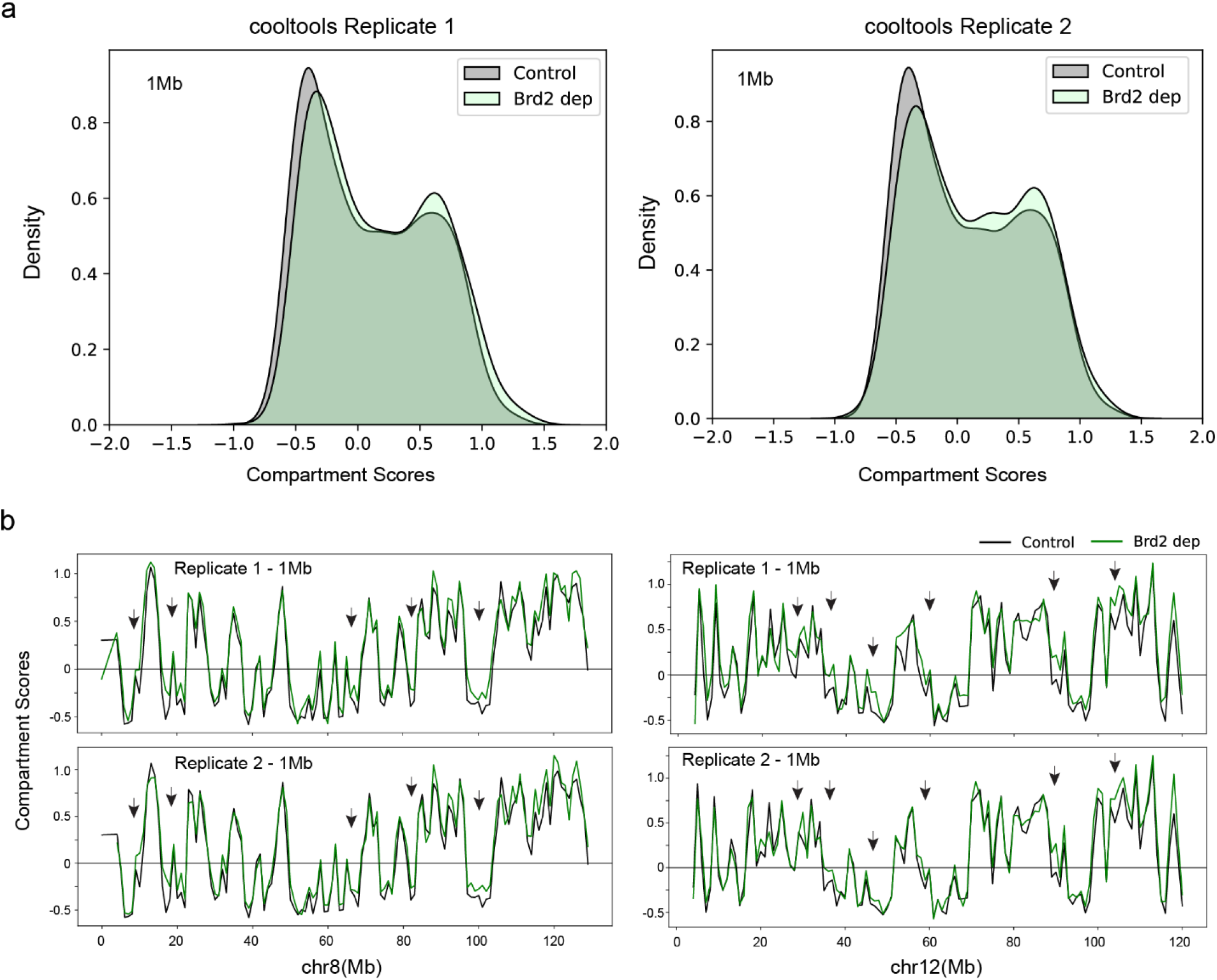
Compartment score distributions for both replicates at 1Mb resolution after correcting the eigenvector selection and orientation inconsistencies (panel a). Compartment score tracks for chr8 and chr12 were shown in panel b. Here, chr8 and chr12 are selected as the two replicates show high concordance between themselves.

### Saddle plots and saddle strengths suggest that Brd2 depletion leads to weakening of compartmentalization

As eigenvectors are only determined up to a scalar multiple, comparing the raw values of the profiles may not be the best metric to differentially analyze contact maps. Saddle plots provide a way to measure the strength of compartmentalization across experiments by using quantiles instead of raw values. Saddle strength plots provide a representation of the ratio between compartment self-interactions (AA + BB) and cross-interactions (AB) across varying quantile widths. These analyses, designed to assess the strength of compartment, suggest that Brd2 depletion leads to a decrease of compartment strength especially among B compartments at multiple resolutions (Figure 3 and supplementary files). This is consistent across chromosomes taken individually as well (Figure 4 and supplementary files).

**Figure 3.**
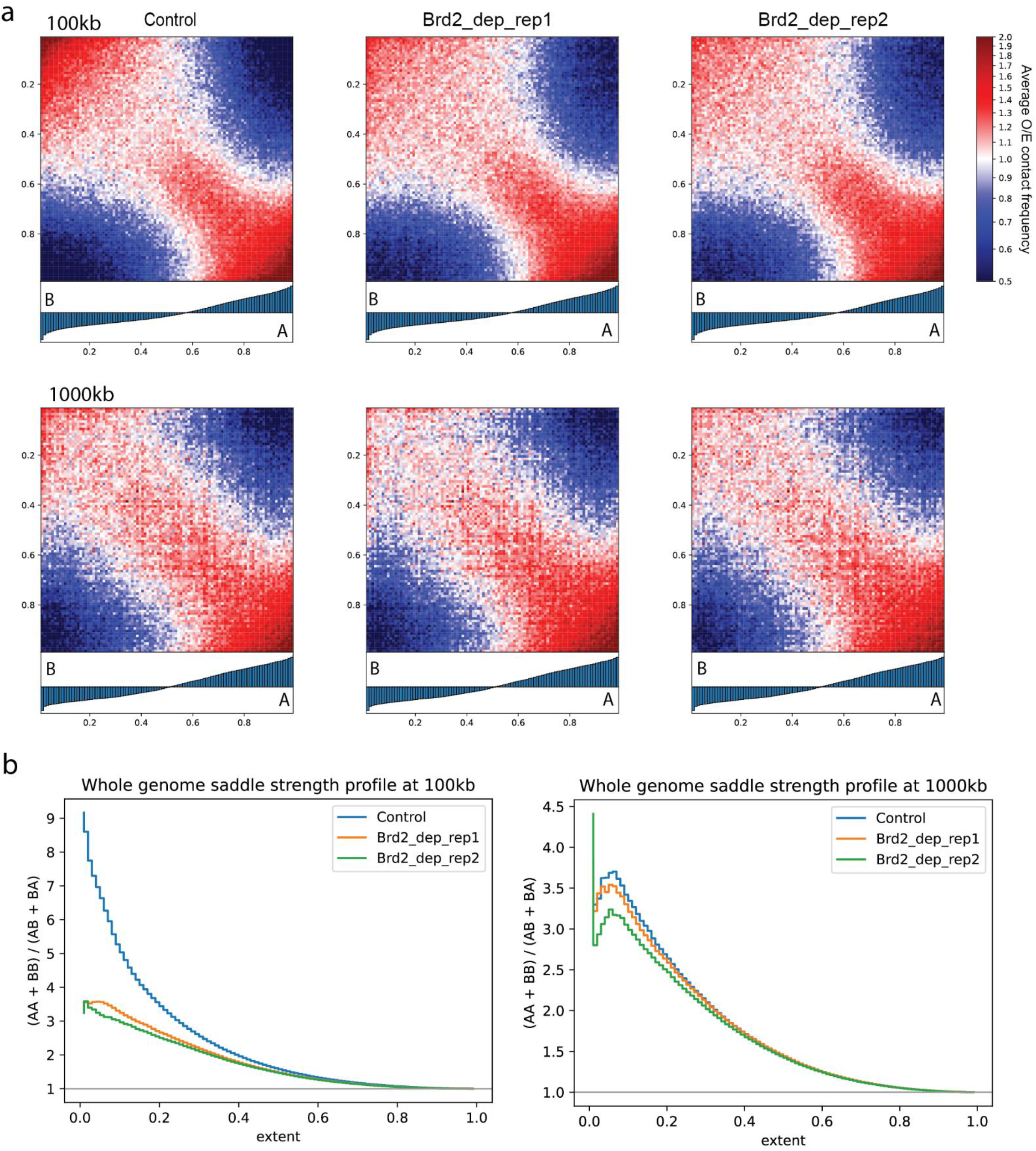
Saddle plots (panel a) and saddle strengths (panel b) for the whole genome, binned into 100 quantiles using the cooltools generated compartment profile for Control at 100kb and 1000kb resolution.

**Figure 4.**
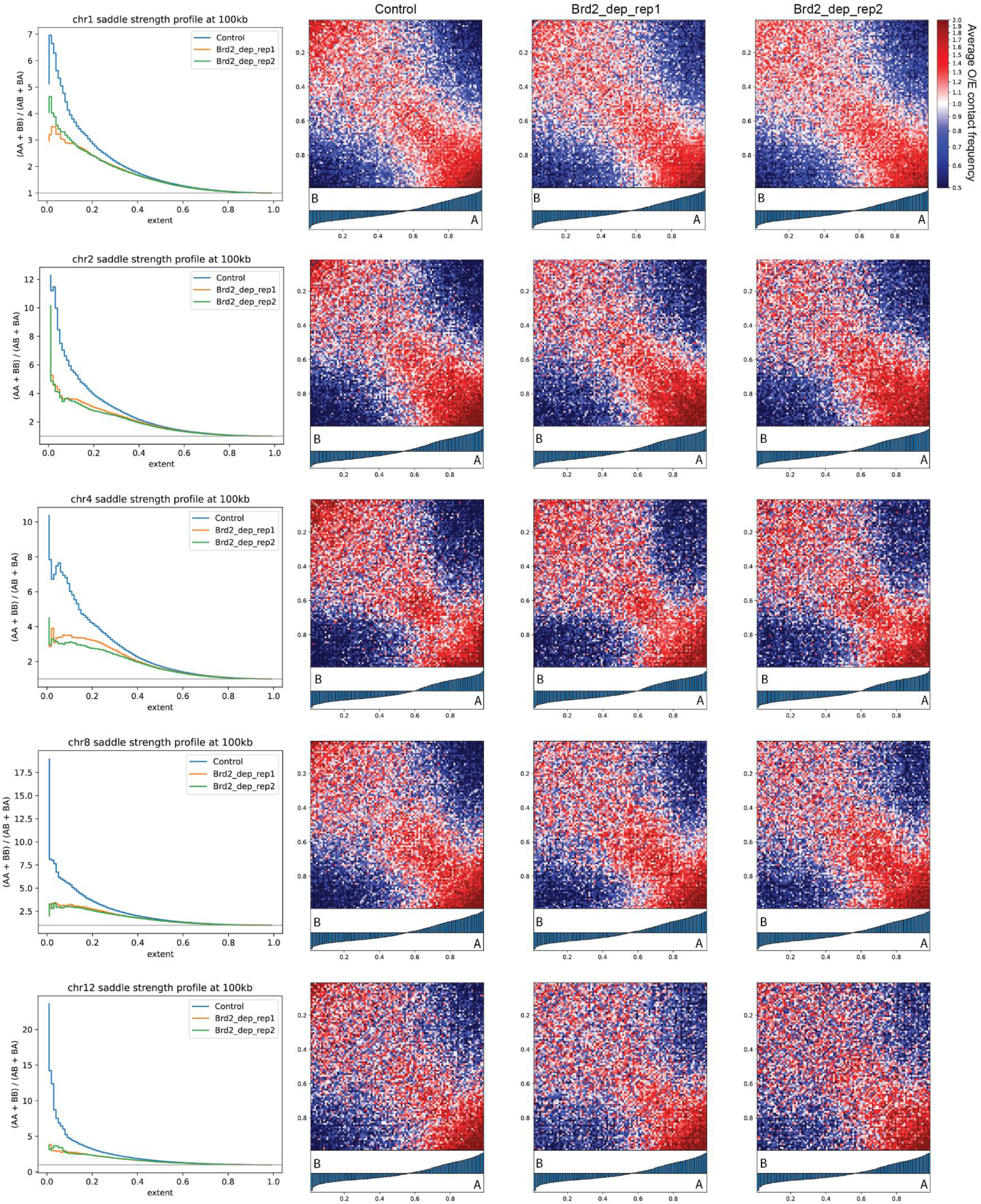
Saddle plots and saddle strengths for chr1, chr2, chr4, chr8 and chr12, binned into 100 quantiles using the cooltools generated compartment profile for Control at 100kb resolution. The weakening of B compartment is evident across individual chromosomes at other resolutions 100kb, 200kb, 500kb and 1Mb (See details in supplementary files).

## Discussion

Our re-examination of the data reveals that the discrepancies between the findings of Hinojosa-Gonzalez et al.^2^ and Xie et al.^1^ likely arise from a combination of different resolutions examined, different computational tools, and inconsistent compartment profile selection and orientation. The study also revealed the susceptibility of compartment profile selection to errors, which can significantly impact data interpretation. This underscores the importance of meticulous manual inspection and selection of appropriate eigenvector, as well as the phase orientation, to ensure accuracy. Comparative analyses should account for both global and local changes at a variety of resolutions, as well as normalization techniques. Moreover, our findings advocate for using saddle and saddle strength plots over compartment score tracks for differential comparisons. Our work emphasizes the need for the development and adoption of standardized, fully automated, and easily understandable computational methodologies. Such standardization is essential for ensuring the consistency and reliability of research findings, which, in turn, will enhance our capacity to derive biologically significant insights from complex genomic data.

## Data and code availability

Datasets used in this study are Control: SRR13291356, Brd2_dep: SRR13291360, submitted to NCBI GEO under the accession number GSE163729. The code used for processing the data and generating the figures, as well as additional supplementary files showing the saddle plots, saddle strengths, compartment profiles, histograms, and scatter plots for multiple resolutions are available online in this GitHub repository: https://github.com/ZhangGroup-MITChemistry/brd2-reanalysis.

## Acknowledgements

We thank Ferhat Ay, David M. Gilbert, Nezar Abdennur, Geoffrey Fudenberg and Laura Hinojosa-Gonzalez for critical comments and feedback on our reanalysis and manuscript.

